# Drivers of diversity within and between microbial communities during stochastic assembly

**DOI:** 10.1101/2024.11.19.624346

**Authors:** Loïc Marrec, Claudia Bank

## Abstract

No two microbial communities share the same species richness and abundance. Experiments have shown that the assembly of new microbial communities from the same environmental pool is sufficient to generate diversity within and between communities. To identify the mechanisms behind these experimental results, we build a stochastic model that considers both the dispersal of microbes from a pool to communities and microbial division. By analyzing timescales, we identify distinct assembly regimes. Specifically, if dispersal is slower than division, microbial communities show low diversity within communities but high diversity between communities. Conversely, if dispersal is faster than division, microbial communities exhibit high diversity within communities but low diversity between them. We validate these predictions both numerically, using Gillespie simulations, and analytically, by deriving equations for species richness and abundance distributions. Our derivations pinpoint two key metrics, the bimodality coefficient and mean relative abundance, which identify the assembly regime and quantify trait differences. We apply these metrics to reanalyze an experimental data set, demonstrating their practical application. Overall, our study provides general predictions on how stochasticity, timescales, and traits impact diversity within and between communities during the assembly of new microbial communities, allowing for a better understanding of variation in microbiome formation.

## Introduction

Microorganisms inhabit marine, terrestrial and host-associated ecosystems. In addition to being everywhere, they are present in large numbers. There are 10^29^ bacteria in the oceans, 10^9^ in a teaspoon of soil, and 10^11^ in a gram of dental plaque (Nat, 2011). Many large microbial communities assemble from scratch when a new micro-environment is formed. For example, volcanic eruptions create sterile environments ripe for new microbial assembly, such as hot springs, fumaroles, and lava tubes (Hadland et al., 2024). Microbial communities play a critical role in numerous biological processes. This sparks great interest from basic and applied science in understanding their structure and function. Recent advances in genomic and metagenomic sequencing have shed light on the coexistence of several microbial species within individual communities, which often build complex multi-species ecosystems. The accumulation of data has shown that no two microbial communities share the same species richness and abundance. The origins and drivers of this vast diversity within and between microbial communities remain to be identified. The growing interest in deciphering the mechanisms underlying such diversity has led to numerous bottom-up experimental studies. Some of them suggest that diversity emerges in the early stages of microbial community assembly (Vega and Gore, 2017; Ortiz et al., 2021; Jones et al., 2022). Comprehensive theoretical models are needed to quantify the processes driving these diversity patterns during microbial community assembly.

Host-microbe associations in the gut are one of the most extensively studied systems due to its relevance for host health and disease (Relman, 2015). Many organisms do not inherit their parent’s microbiota and, thus, are born with germ-free guts that are populated after birth [e.g., *C. elegans* (Zhang et al., 2017), *D. melanogaster* (Blum et al., 2013)], making community assembly a potentially important stage in the emergence of diversity. Other organisms, like humans, inherit a small fraction of their parents’ gut microbiota (Perez-Muñoz et al., 2017). In the gut of animal hosts, host health is crucially impacted by microbial communities that facilitate food digestion or cause diseases (Gilbert et al., 2018). For instance, humans with Crohn’s disease have a lower microbial diversity and different composition than healthy humans (Manor et al., 2020), suggesting that high diversity in host-associated microbial communities is related to good host health (Lozupone et al., 2012; Mosca et al., 2016; Gilbert et al., 2018). In addition to diversity within individual communities, microbial communities also exhibit high diversity between communities. For example, substantial variation in microbial community structure (i.e., composition and abundances) is observed among hosts of the same species (Spor et al., 2011; Lundberg et al., 2012; Smith et al., 2015). Even monozygotic twins were shown to have high within-twin-pair microbiome diversity (Goodrich et al., 2014; Vilchez-Vargas et al., 2022).

To identify the key drivers of diversity within and between microbial communities, experimental studies increasingly use a *bottom-up* approach to assemble new communities. Assembling microbial communities under controlled conditions, such that experimentalists can systematically manipulate factors (e.g., species composition), helps quantify how these factors contribute to the emergence and maintenance of diversity within individual communities. Additionally, when scaled across multiple microbial communities, these experiments provide insights into how variations in microbial community structure arise. Such bottom-up experiments are essential to isolating the mechanisms that govern diversity patterns, advancing our understanding of microbial ecology. For example, Vega and Gore (2017) carried out an experiment involving initially germ-free *C. elegans* worms fed on *E. coli* bacteria. They showed that community assembly in the intestines of hosts that are isogenic, isolated, and fed on the same bacterial pool, produces substantial inter-host heterogeneity (Vega and Gore, 2017). Their results highlighted that the early stages of microbial communities are not only driven by deterministic mechanisms, such as selection, but also by stochasticity. However, the importance of random events in shaping microbial communities relative to deterministic mechanisms is contentious (Zhou and Ning, 2017). There is therefore a strong need for mathematical models accounting for stochasticity to better understand diversity patterns.

Here, we build a stochastic model inspired by previous work from Houchmandzadeh (2018) and Vega and Gore (2017), which accounts for two microbial features/traits: the rate of microbial dispersal from a pool to communities and the division rate. By analyzing timescales of dispersal and division, we identify distinct assembly regimes leading to different diversity patterns. We validate these predictions with numerical simulations and analytical derivations for species richness and abundance distributions. Our derivations indicate two key metrics, the bimodality coefficient and the mean relative abundance, which determine the assembly regime and quantify trait differences. To demonstrate their practical application, we apply these metrics to experimental data from Vega and Gore (2017). Overall, our study quantifies how stochasticity, timescales, and traits impact diversity within and between communities during the assembly of new microbial communities, allowing for a better understanding of microbiome formation.

## Model and methods

### Continent-island model of community assembly

We represent the assembly of microbial communities using a continent-island model (Latter, 1973). In the main text, we focus on two-species communities; we relax this assumption in the Supplementary Material, Section 4. Each microbial species is described by two “trait” parameters that account for processes known to crucially impact community assembly: dispersal and division (Vellend, 2010; Costello et al., 2012). As shown in Figure 1, microbes disperse from the microbial pool (i.e., the continent; also called regional species pool in ecology) into the communities (i.e., the islands). The microbial pool is fixed and composed of two species A and B, whose division rates are denoted by *r*_A_ and *r*_B_, respectively. The sign of the selection coefficient *s*, which is given by *s* = *r*_A_ − *r*_B_, indicates whether A microbes are favored (i.e., *s >* 0) or disfavored (i.e., *s <* 0) by natural selection within a community.

**Figure 1:**
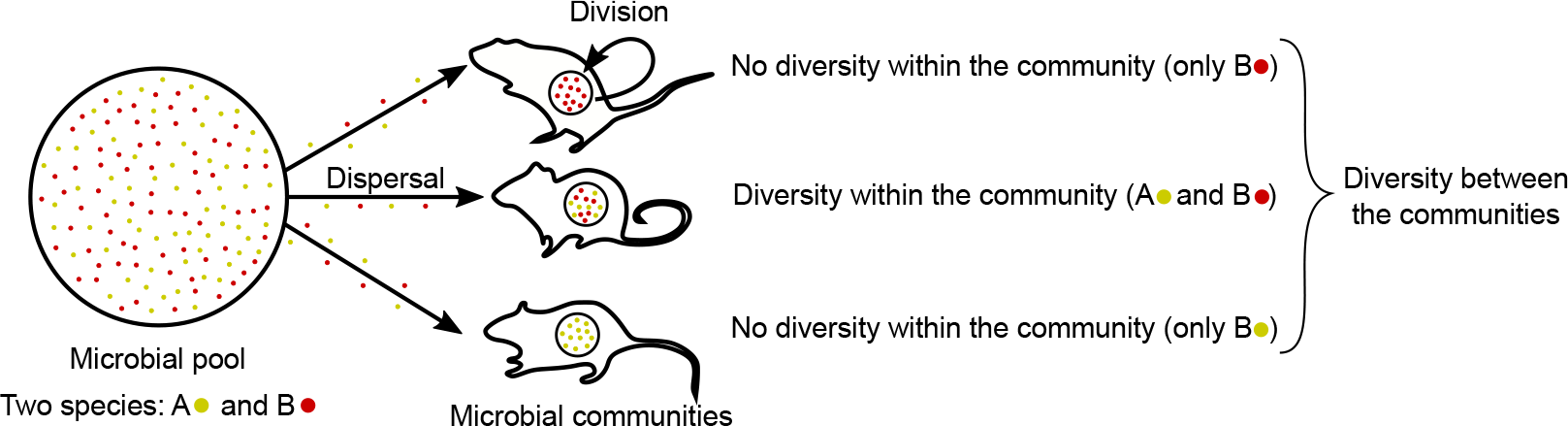
Illustration of the model - Microbial community assembly. The environmental microbial pool contains two species, A (yellow) and B (red). Microbes disperse from the pool into local microbial communities and divide within these communities. Once the carrying capacity of the microbial community is reached, we determine whether the two species coexist (diversity within the community) or do not coexist (no diversity within the community). We compare the composition of several communities to quantify diversity between communities.

The two species may be present at unequal abundances in the microbial pool or have different abilities to populate a community. To account for this, we introduce species-dependent dispersal rates, i.e., *c*_A_ and *c*_B_. The size of each community, denoted by *N* = *N*_*A*_ + *N*_*B*_, can increase over time but is upper-bounded by a constant carrying capacity (i.e., 0 ≤ *N* ≤ *K*). This carrying capacity may result, for example, from limited space. The microbial communities follow a logistic growth such that the *per capita* division rate of species A microbes is *r*_A_(1 − *N/K*) and the *per capita* division rate of species B microbes is *r*_B_(1 − *N/K*). Note that our approach is extendable to other types of growth [e.g., Gompertz, Richards, etc; see Tsoularis and Wallace (2002)] by changing the form of density dependence.

### Gillespie algorithm simulating the microbial community assembly

We simulate the assembly of a microbial community with a Gillespie algorithm (Gillespie, 1976, 1977). This algorithm generates trajectories of stochastic dynamics whose event rates are known. Our model features four types of events: the divisions of the microbes of species A and B

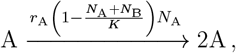

and

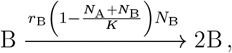

and the dispersal of the microbes of species A and B

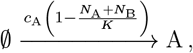

and

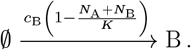

For each event, the propensity function is indicated above the arrow. The simulation steps are as follows:

1. Initialization: The microbial community starts from *N*_A_ = 0 and *N*_B_ = 0 microbe at time *t* = 0.
2. Time update: The time increment Δ*t* is randomly sampled from an exponential distribution with mean 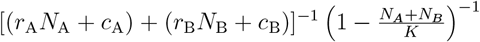 and the time is updated by *t* ← *t* + Δ*t*.
3. Event selection: The next event to occur is chosen randomly proportionally to its probability. For example, the division of a microbe of species A is chosen with probability *r*_A_*N*_A_*/*[(*r*_A_*N*_A_ + *c*_A_) + (*r*_B_*N*_B_ + *c*_B_)].
4. Population size update: The population sizes are updated according to the event selected in Step 3. For example, if the division of a species A microbe is chosen, the population size of species A is updated by *N*_A_ ← *N*_A_ + 1.
5. We return to Step 2 until the microbial community is populated up to the carrying capacity (i.e., *N*_A_ + *N*_B_ = *K*).

In the following, we consider that each stochastic realization of the above algorithm describes the assembly of a single microbial community. Thus, collecting several stochastic realizations is equivalent to simulating the microbial assembly of several communities.

### Data availability

Simulations were performed with Matlab (version R2021a). All annotated code to reproduce the simulations and visualizations is available at https://github.com/LcMrc and will be deposited on Zenodo upon acceptance of the paper.

## Results

### The relationship between dispersal and division drives the assembly of microbial communities

In our model, a new community is assembled by microbes from a microbial pool in the environment. In Vega and Gore (2017)’s experiment, in which worms feed on bacteria, the dispersal rate depends on the bacterial densities in the microbial pool. A high density of microbes induces a high dispersal rate and vice versa. Once inside the community, microbes divide and compete, populating the microbiota. The community assembly is thus composed of two processes: dispersal and division. For simplicity, we initially consider a neutral case in which the two microbial species have the same division rate and dispersal rate (i.e., *r*_A_ = *r*_B_ = *r* and *c*_A_ = *c*_B_ = *c*). The time between two dispersal events of microbes that establish in the microbiota, denoted by *T*_*c*_, is equal to the inverse of the dispersal rate, i.e. *T*_*c*_ = 1*/c*. The growth time of a microbial community starting from an individual until the carrying capacity is reached, assumed without any microbial dispersal, is given by 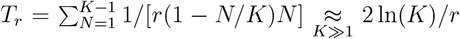. This expression for the growth time assumes logistic growth and is obtained by summing the average of each division time. Both timescales are equal if and only if *T*_*c*_ = *T*_*r*_. Defining *c*_lim_ as the dispersal rate for which this equality is satisfied, we obtain

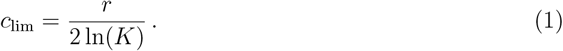

Thus, as in Vega and Gore (2017)’s work, we expect community assembly to be driven by microbial dispersal if *c > c*_lim_, by microbial dispersal and division if *c* ∼ *c*_lim_, and by microbial divisions if *c < c*_lim_. However, since Vega and Gore (2017) focused on the exponential growth phase, they found *c*_lim_ = *r*.

### A stochastic model predicts species richness and abundance

#### Master equation

The assembly of a new microbial community is a stochastic process that involves, in our model, two types of events: dispersal of a microbe from the microbial pool to the community, and microbial division. As the microbial community is assumed to be initially microbe-free, every assembly process includes a phase in which the community size is small. During this phase, stochasticity plays a major role, regardless of the carrying capacity. We, therefore, use a microscopic and probabilistic description of the microbial community assembly similar to the model developed by Houchmandzadeh (2018), which includes dispersal in addition to division. Our model is similar to that of Vega and Gore (2017), neglecting death for mathematical convenience; we relax this assumption in the Supplementary Material, Section 5. Specifically, we analyze a system of equations governing the dynamics of the probability *P* (*N*_A_, *N*) of having *N*_A_ microbes of species A in a community of size *N* (and thus *N*_B_ = *N* −*N*_A_ microbes of species B). This system of equations, called the master equation (Gardiner, 2009; Van Kampen, 2011), satisfies

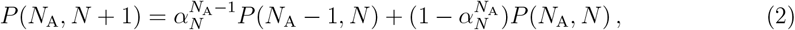

where 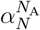 is the probability that the increase in community size by one individual (i.e., *N* →*N* + 1) is due to the dispersal or division of a microbe of species A. This probability is given by

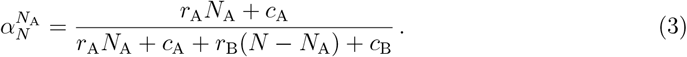

As pointed out by Houchmandzadeh (2018), this probability does not depend on the type of density dependence. It is thus valid for other growth types [e.g., Gompertz, Richards, etc; see Tsoularis and Wallace (2002)]. Since we assume that the microbial assembly starts from a microbe-free community, the master equation 2 admits as initial condition *P* (0, 0) = 1. Moreover, at the two boundaries of the master equation 2 (i.e., for *N* = 0 and *N* = *K*), it reduces to a one-term recurrence relation. For example, considering the lower boundary *N* = 0 leads to a recurrence relation on the probability *P* (0, *N*) of having no microbe of species A in a community of size *N*, which reads

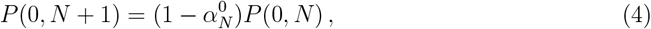

the solution of which is

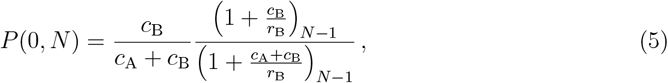

where (*x*)_*N*_ = *x*(*x* + 1)(*x* + 2)…(*x* + *N* − 1) is the Pochhammer symbol. Another way to recover Equation 5 is to note that 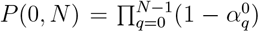. Similarly, we get an expression for the probability *P* (*N, N*) of having only microbes of species A in a community of size *N*, noting that 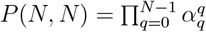, which leads to

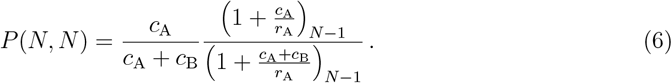

By evaluating Equations 5 and 6 at *N* = *K*, i.e., *P*_0_ = *P* (0, *K*) and *P*_*K*_ = *P* (*K, K*), respectively, we determine whether coexistence of both microbial species occurs at the end of the assembly process. More specifically, the probability that both microbial species coexist, denoted by *P*_coexist_, is given by

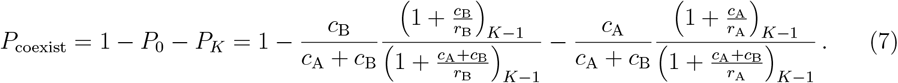

#### Low dispersal regime

When the dispersal rates are much lower than the division rates (i.e., *c*_A_, *c*_B_ ≪ *r*_A_, *r*_B_), the probability 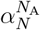 (see Equation 3) simplifies to

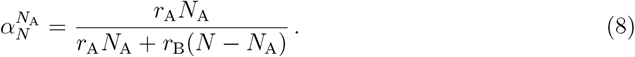

The resulting master equation was extensively investigated by Houchmandzadeh (2018) (see Equation 2). It cannot be analytically solved if both division rates differ, although some approximations exist [see Houchmandzadeh (2018) for more detail]. Note, however, that Equations 5 and 6 simplifies to

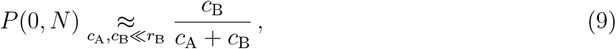

and

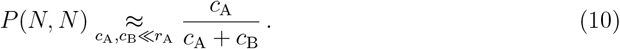

when the dispersal rates are much lower than the division rates.

#### High dispersal regime

When the dispersal rates are much larger than the division rates (i.e., *c*_A_, *c*_B_ ≫ *r*_A_, *r*_B_), the probability 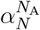 (see Equation 3) simplifies to

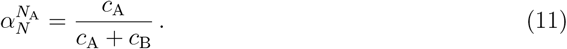

This simplification allows us to solve the master equation (see Equation 2), the solution of which reads

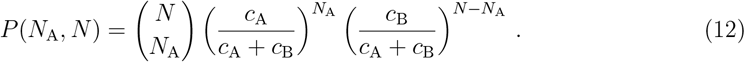

Thus, the number of A microbes follows a binomial distribution in the high dispersal regime.

#### Neutral case

Here, we assume that both microbial species, namely A and B, have the same dispersal and division rates. This neutral case was experimentally investigated by Vega and Gore (2017) using two bacterial strains that only differed in their fluorescent label (YFP and dsRed). Such neutrally competing microbes should satisfy *r*_A_ = *r*_B_ = *r* and *c*_A_ = *c*_B_ = *c*, which simplifies the probability 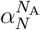 to

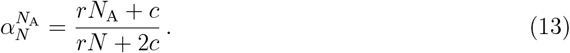

Here, the probability 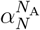 becomes linear in *N*_A_. This enables us to derive the n^th^ moment of *N*_A_(*N*), i.e.,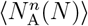, which denotes the number of A microbes when the community size equals *N*. In particular, considering 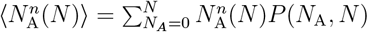 (see Equation 2) leads to

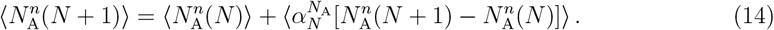

The first four moments, which describe the probability distribution of the stochastic variable *N*_A_(*N*), are given by

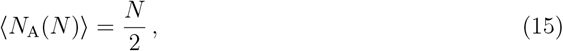

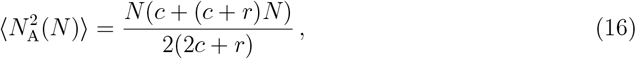

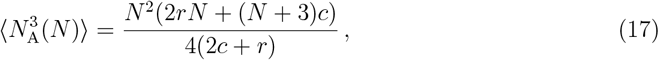

and

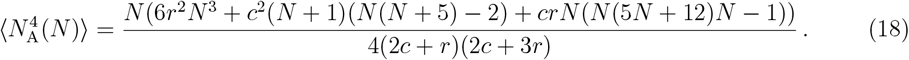

The variance *σ*^2^ reads

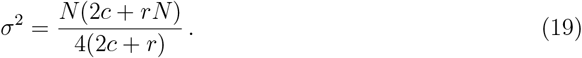

Note that assuming *r* ≫ *c* and *N* ≫ 1 leads to ⟨*N*_A_(*N*)⟩^2^ = *σ*^2^, revealing giant fluctuations in the number of microbes (Houchmandzadeh, 2018). The skewness of the distribution is zero, i.e., *γ* = 0, as expected for neutrally assembling microbes. The kurtosis *κ* reads

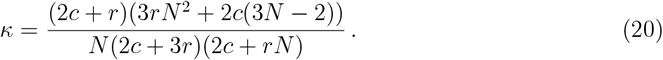

From the variance *σ*^2^ and the kurtosis *κ*, we derive the bimodality coefficient (Ellison, 1987), denoted by BC, which, according to Sarl’s definition, is given by

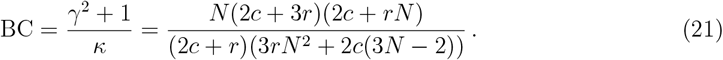

Note that the bimodality coefficient was experimentally measured by Vega and Gore (2017) to relate the heterogeneity of microbiota to the microbial density present in the pool from which the hosts fed. The BC is a summary statistic of probability distributions that ranges from 0 to 1. A bimodality coefficient equal to 1 corresponds to a Bernoulli distribution, whereas a bimodality coefficient equal to 5/9 corresponds to a uniform distribution. Thus, BC values greater than 5/9 indicate a bimodal distribution, whereas BC values lower than 5/9 indicate an unimodal distribution. Assuming that the dispersal rate is much lower than the division rate (i.e., *c* ≪ *r*) leads to a bimodality coefficient equal to 1, resulting in a Bernoulli distribution where the final number of A microbes is zero with probability 1/2 and *N* with probability 1/2 (see Equations 9 and 10 with *c*_A_ = *c*_B_). Conversely, assuming the opposite (i.e., *c* ≫ *r*) leads to a bimodality coefficient equal to 1/3, resulting in a binomial distribution satisfying Equation 12. Finally, assuming that the division and dispersal rates are equal (i.e., *r* = *c*), in addition to assuming that *N* ≫ 1, yields a bimodality coefficient equal to 5/9, resulting in a uniform distribution satisfying Equation 23 (see paragraph below).

#### Neutral case - Intermediate dispersal regime

When the dispersal and division rates are equal (i.e., *c*_A_ = *c*_B_ = *r*_A_ = *r*_B_), the probability 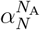 simplifies into

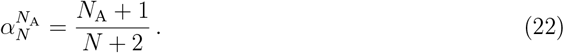

Here, the probability satisfying

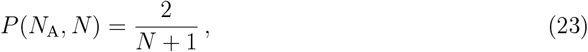

is a solution of the master equation 2. Here, *P* (*N*_A_, *N*) is independent of the number of A microbes, and the structure of microbial communities follows a uniform distribution.

#### Deterministic predictions

In the Supplementary Material, Section 1, we derive analytical predictions using a deterministic model accounting for the same events as its stochastic counterpart. We use these predictions to quantify the species abundances and compare them to the predictions obtained with the stochastic model.

### Dispersal and division timescales shape diversity within and between communities

We first focus on two microbial species that neutrally compete. In other words, both microbial species have the same dispersal and division rates (i.e., *c*_A_ = *c*_B_ = *c* and *r*_A_ = *r*_B_ = *r*), which allows us to assess the impact of dispersal and division timescales on community assembly. More specifically, we consider the cases in which microbes divide more often than disperse (i.e., *r > c*), divide as often as disperse (i.e., *r* = *c*), and divide less often than disperse (i.e., *r < c*). Figures 2A, B, and C show that these three cases lead to different outcomes for the diversity within and between communities.

**Figure 2:**
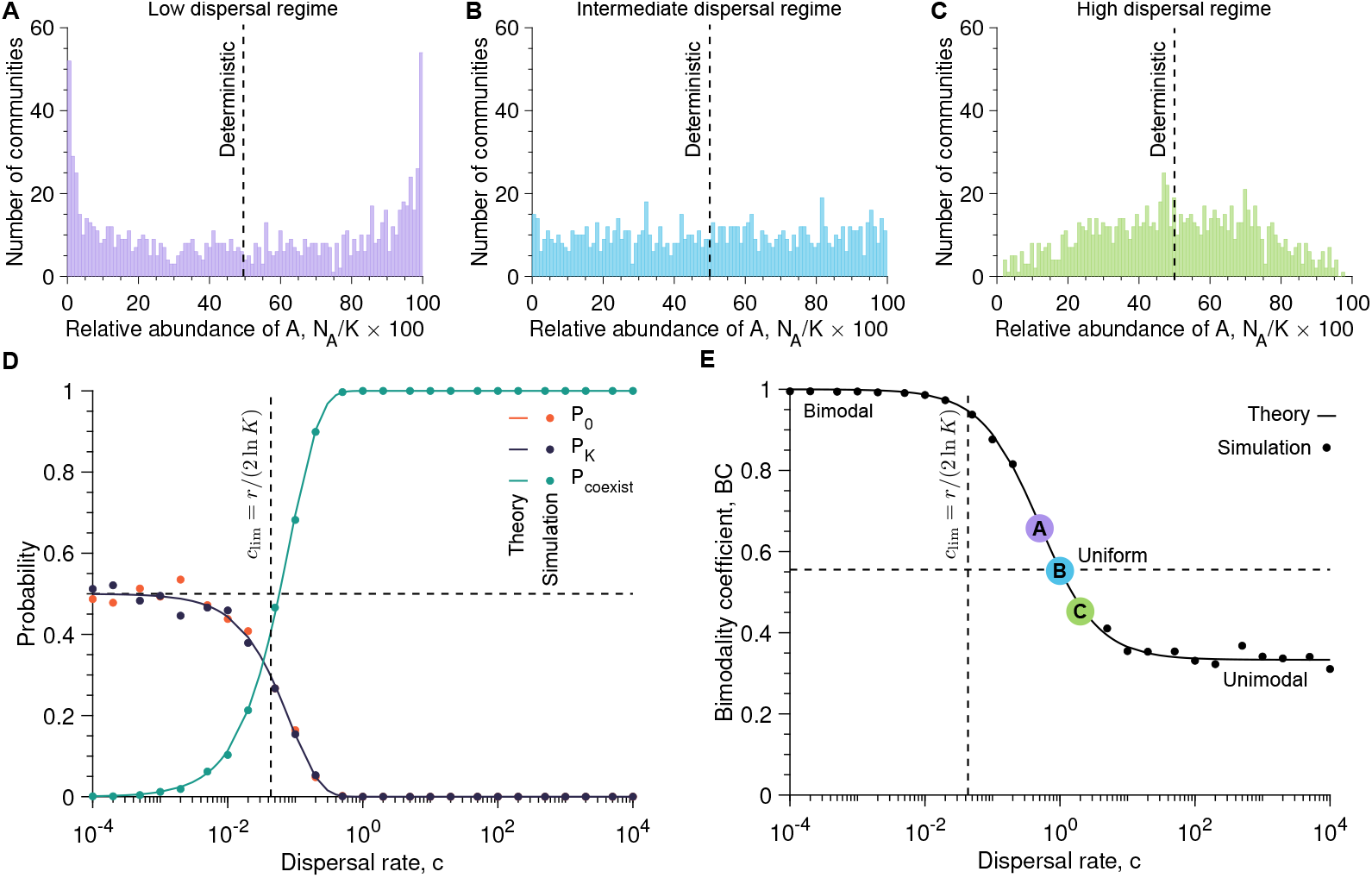
Microbial community assembly depends on how quickly new microbes disperse and how fast they divide once in the community. Sub-figures **A, B**, and **C** show the number of communities versus the relative abundance of A microbes once the carrying capacity is reached (i.e., *N*_A_*/K* × 100) with the dispersal rate equal to *c* = 0.5, *c* = 1, and *c* = 2, respectively (here *c*_A_ = *c*_B_ = *c*). Sub-figure **D** represents the probabilities that, at the end of the microbial community assembly, the community is left with no A microbes (i.e., *P*_0_), has only A microbes (i.e., *P*_*K*_), and is composed of both species (i.e., *P*_coexist_ = 1 − *P*_0_ − *P*_*K*_) as a function of the dispersal rate *c*. Sub-figure **E** presents the bimodality coefficient BC against the dispersal rate *c*. In Sub-figures **D** and **E**, the solid lines show analytical predictions (see Equations 5, 6, 7, and 21), whereas the markers correspond to simulated data averaged over 10^3^ stochastic replicates, each considered a community. In Sub-figures **D** and **E**, the vertical dashed line shows the dispersal rate value at which the timescales associated with dispersal and division are equal (i.e., *c* = *c*_lim_; see Equation 1). In Sub-figures **D** and **E**, the vertical dashed line shows a probability of 1/2 and a bimodality coefficient of 5/9, respectively. Parameter values: division rates *r*_A_ = *r*_B_ = 1, carrying capacity *K* = 10^5^, number of communities 10^3^.

If microbes divide more often than disperse (i.e., *r > c*), the abundance of A microbes in individual communities follows a bimodal distribution (i.e., with two peaks; see Figure 2A). The first microbe dispersing into the community divides and populates it before a second dispersal event occurs. Since A and B microbes have the same dispersal rate, both peaks in relative abundance observed in Figure 2 have the same amplitude. In the extreme case in which microbes divide much more often than disperse (i.e., *r* ≫ *c*), the number of A microbes follows a Bernoulli distribution (see Equations 9 and 10 and Figure S2), such that half of the communities are populated with only A microbes whereas half of the communities are populated with only B microbes. Therefore, the *low dispersal regime* induces low diversity within communities but high diversity between communities (i.e., low *α*-diversity but high *β*-diversity; see Figures S7 and S8).

If microbes divide as often as disperse (i.e., *r* = *c*), the abundance of A microbes in individual communities follows a uniform distribution (see Figure 2B). Both division and dispersal contribute to the community assembly, resulting in numerous possible microbial structures, which are all achieved with the same probability (see Equation 23). Therefore, the *intermediate dispersal regime* induces high diversity within and between communities (i.e., high *α*- and *β*-diversities; see Figures S7 and S8).

If microbes divide less often than disperse (i.e., *r < c*), the abundance of A microbes in individual communities follows a unimodal distribution (i.e., with a single peak; see Figure 2C). The microbial communities are mostly populated with dispersing microbes before any division occurs. In the extreme case in which microbes divide much less often than disperse (i.e., *r* ≪ *c*), the abundance of A microbes follows a binomial distribution (see Equation 12 and Figure S2-B). This binomial distribution shows that the microbial community structure in each community reflects the dispersal rates of A and B microbes, which are here equal, thus resulting in a single peak around 50% of microbes A, and thus 50% of microbes B, in individual communities (see Figure 2C). Therefore, the *high dispersal regime* induces low diversity between communities but high diversity within communities (i.e., high *α*-diversity but low *β*-diversity; see Figures S7 and S8).

As reported in Figures 2D and E, the probability of coexistence of both microbial species within a community *P*_coexist_ and the bimodality coefficient BC illustrate the transition from non-coexistence, observed when microbes divide more often than disperse (i.e., *r > c*), to coexistence, observed when microbes divide less often than disperse (i.e., *r < c*). On the one hand, as reported by Figure 2D, the probability of coexistence *P*_coexist_ increases from 0 to 1 as the dispersal rate *c* increases, passing through 1/2 when the dispersal rate satisfies *c* = *c*_lim_ (see Equation 1), i.e., when both dispersal and division timescales are equal. Our analytical predictions, given by Equations 5, 6, and 7 allow us to predict whether both microbial species coexist at the end of the community assembly and, thus, whether high diversity within communities emerges.

On the other hand, as shown by Figure 2E, the bimodality coefficient BC decreases from 1, corresponding to a bimodal Bernoulli distribution and synonymous with low diversity within hosts and high diversity between communities (i.e., low *α* diversity but high *β* diversity), to 1/3, corresponding to a unimodal distribution and synonymous with high diversity within communities and low diversity between communities (i.e., high *α* diversity but low *β* diversity), as the dispersal rate *c* increases. Therefore, in addition to predicting whether high diversity within and between communities likely occurs or not, Equation 21 quantifies the exact shape of the species abundance distribution describing the microbial community structure.

Our results highlight how a stochastic formalism is necessary to capture the fluctuations in the abundance of A and B microbes during assembly of a new community and, therefore, to characterize diversity within and between communities. Both the deterministic and stochastic approaches predict that, on average, neutrally competing species will be present in the community at equal abundances (i.e., *N*_A_ = *N*_B_ = *K/*2; see Equations S5 and 15). Although our simulated data validates this prediction, it does not reflect the observed microbial community structure. In particular, Figures 2A, B, and C show that, for every dispersal rate, the mean percentage of A microbes at the end of the community assembly is 50%. However, Figure 2A reveals that very few communities are populated with 50% of A and B microbes when the dispersal rate is low. Conversely, Figure 2C shows a high number of communities with 50% of A and B microbes for higher dispersal rates. The initial stochastic period in the low dispersal regime thus crucially influences the observed abundance distributions independent of the final community size.

### Differences in dispersal rates lead to asymmetry in species abundances

Next, we assume that the two microbial species have the same division rates but different dispersal rates (i.e., *r*_A_ = *r*_B_ and *c*_A_ ?= *c*_B_). As a reminder, differences in dispersal rates may result from a different relative abundance in the microbial pool or a different ability to persist in the microbial community after dispersal. Figure 3 illustrates a case in which A microbes disperse twice as often as B microbes (i.e., *c*_A_ = 2*c*_B_). Similar to the neutral case, Figure 3 shows a transition from a bimodal distribution of the relative abundance of A microbes in the microbial communities, obtained for low dispersal rates compared to division rates (i.e., *r > c*), to an unimodal distribution, obtained for high dispersal rates compared to division rates (i.e., *r < c*). However, as opposed to the neutral case, the distribution of the relative abundance of A microbes in the microbial communities is no longer symmetric.

**Figure 3:**
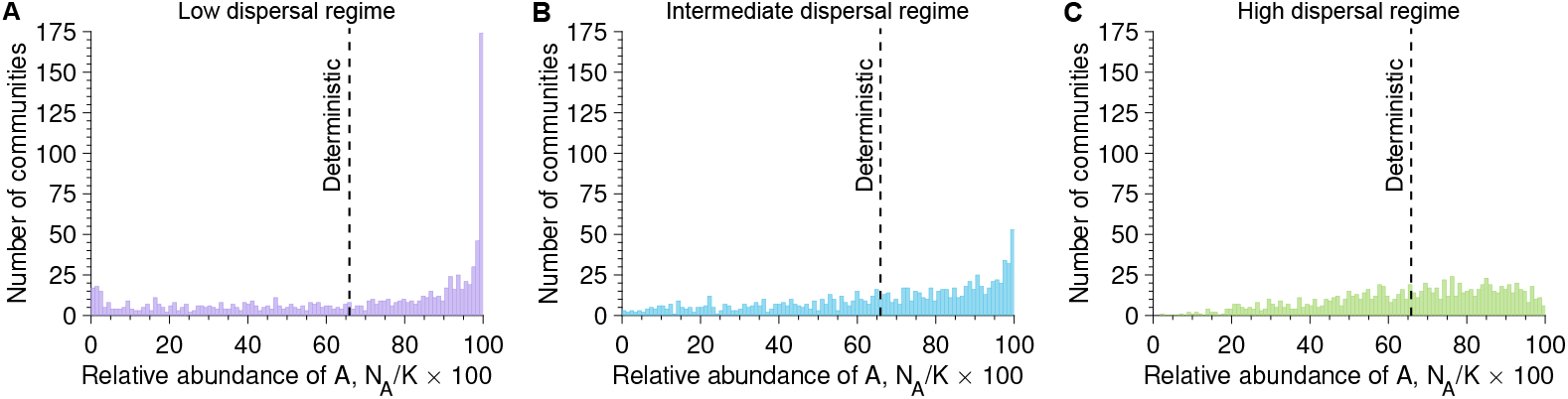
Differences in dispersal rates induce asymmetric species abundance distributions. Sub-figures **A, B**, and **C** show the number of communities versus the relative abundance of A microbes once the carrying capacity is reached (i.e., *N*_A_*/K* × 100) with the dispersal rate equal to *c* = 0.5, *c* = 1, and *c* = 2, respectively [here *c*_A_ = 2*c*_B_ and *c* = (*c*_A_ + *c*_B_)*/*2]. Parameter values: division rates *r*_A_ = *r*_B_ = 1, carrying capacity *K* = 10^5^, number of communities 10^3^.

In the low dispersal regime, Figure 3A shows that a majority of communities are now populated with only A microbes. In this regime, community assembly is driven by the first microbe dispersing from the microbial pool, which is, here, twice as probable to be an A microbe than a B microbe as *c*_A_ ≠ 2*c*_B_. When dispersal rates are extremely low compared to division rates (i.e., *c*_A_, *c*_B_ ≪ *r* with *c*_A_ = 2*c*_B_ and *r*_A_ = *r*_B_ = *r*), a fraction of *c*_A_*/*(*c*_A_ + *c*_B_) communities is populated with only A microbes, whereas a fraction of *c*_B_*/*(*c*_A_ +*c*_B_) communities are populated only with B microbes (see Equations 9 and 10 and Figure S3-A). Therefore, similar to the neutral case, the low dispersal regime leads to no diversity within communities but high diversity between communities (i.e., low *α* diversity but high *β* diversity).

In the high dispersal regime, Figure 3C shows that most communities are populated with a majority of A microbes. In this regime, community assembly is mostly driven by dispersal events rather than divisions. As A microbes disperse more frequently than B microbes (*c*_A_ = 2*c*_B_ in Figure 3B), the peak in the distribution of the relative abundance of A microbes is shifted from the center for the neutral case (i.e., 50%) to the right (i.e., between 50% and 100%). In the extreme case in which microbes divide much less often than disperse (i.e., *c*_A_, *c*_B_ ≫ *r*), most hosts are populated with *c*_A_*K/*(*c*_A_ + *c*_B_) A microbes and *c*_B_*K/*(*c*_A_ + *c*_B_) B microbes (see Equation 12 and Figure S3-B). Therefore, similar to the neutral case, the high dispersal regime leads to high diversity within communities but low diversity between communities (i.e., high *α* diversity but low *β* diversity).

### Differences in division rates become invisible in the very low and high dispersal limits

Next, we assume that the two microbial species have the same dispersal rates but different division rates, resulting in nonzero selection coefficients (i.e., *c*_A_ = *c*_B_ and *r*_A_ ≠ *r*_B_, leading to *s*≠ 0).

Figure 4 shows simulated data for a selection coefficient of *s* = 0.05. In this case, A microbes divide slightly more often than B microbes, giving them a division advantage. Independent of whether microbes divide more often than they disperse (i.e., *r*_A_, *r*_B_ *> c*), as often (i.e., *r*_A_, *r*_B_ ≈ *c*), or more often (i.e., *r*_A_, *r*_B_ *> c*), division advantage induces an asymmetry in the distribution of the relative abundance of A microbes. More specifically, we observe that most microbial communities are composed of a majority of A microbes. Therefore, asymmetry in the relative abundance of A microbes can reflect either the presence of species-specific division or dispersal or advantage.

**Figure 4:**
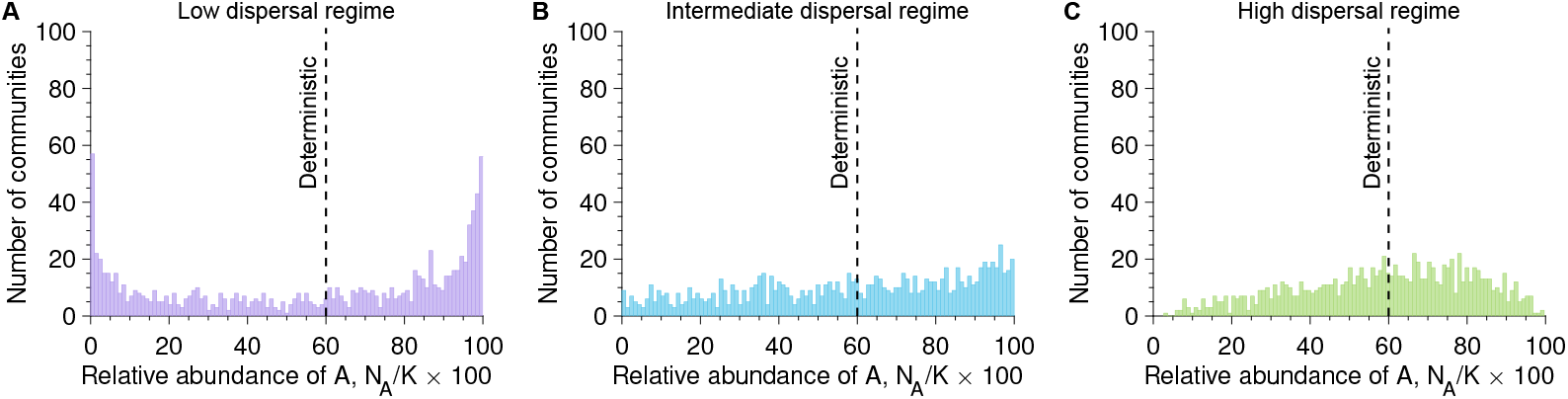
Differences in division rates do not always induce asymmetric species abundance distributions, as opposed to differences in dispersal rates. Sub-figures **A, B**, and **C** show the number of communities versus the relative abundance of A microbes once the carrying capacity is reached (i.e., *N*_A_*/K* × 100) with the dispersal rate equal to *c* = 0.5, *c* = 1, and *c* = 2, respectively (here *c*_A_ = *c*_B_). Parameter values: division rate of A microbes *r*_A_ = 1.05, division rate of B microbes *r*_B_ = 1, selection coefficient *s* = 0.05, carrying capacity *K* = 10^5^, number of communities 10^3^.

Only in the limits of very low and very high dispersal rates, division advantage plays no role in the assembly of microbial communities. As a reminder, the assembly in the very low dispersal regime is driven by the first dispersing microbe, whereas the very high dispersal regime is driven only by dispersal events. Thus, as shown in Figure S4, the distributions of the relative abundance of A microbes do not reflect the differences in division rates and are the same as in the neutral case. In other words, from communities that were assembled at very low or very high dispersal rates, it is impossible to infer whether species that seed the community are subject to within-community selection.

### The bimodality coefficient and mean relative abundance help interpret experimental data

Owing to the stochastic model, we identified two crucial metrics that characterize the abundance distributions: the bimodality coefficient (or the number of peaks in relative abundance distributions) and the mean relative abundance of a microbial species. Here, we argue that these metrics could be used to infer dispersal and division rates from microbial community data. Intuitively, the bimodality coefficient allows for assessing how the dispersal and division timescales compare. If the relative abundance distribution has several peaks, the dispersal rates are much lower than the division rates (i.e., *r*_A_, *r*_B_ ≫ *c*_A_, *c*_B_). In contrast, if the distribution has one peak, the dispersal rates are much larger than the division rates (i.e., *r*_A_, *r*_B_ ≪ *c*_A_, *c*_B_). Second, asymmetric relative abundance distributions reflect differences in dispersal or division rates. To determine which of both drives asymmetry requires data across different dispersal rates. The signature of differences in division rates is expected to disappear at very high and very low dispersal rates, whereas the signature of differences in dispersal rates remains visible at any dispersal rate value. Figure 5 provides a summary of our model predictions.

**Figure 5:**
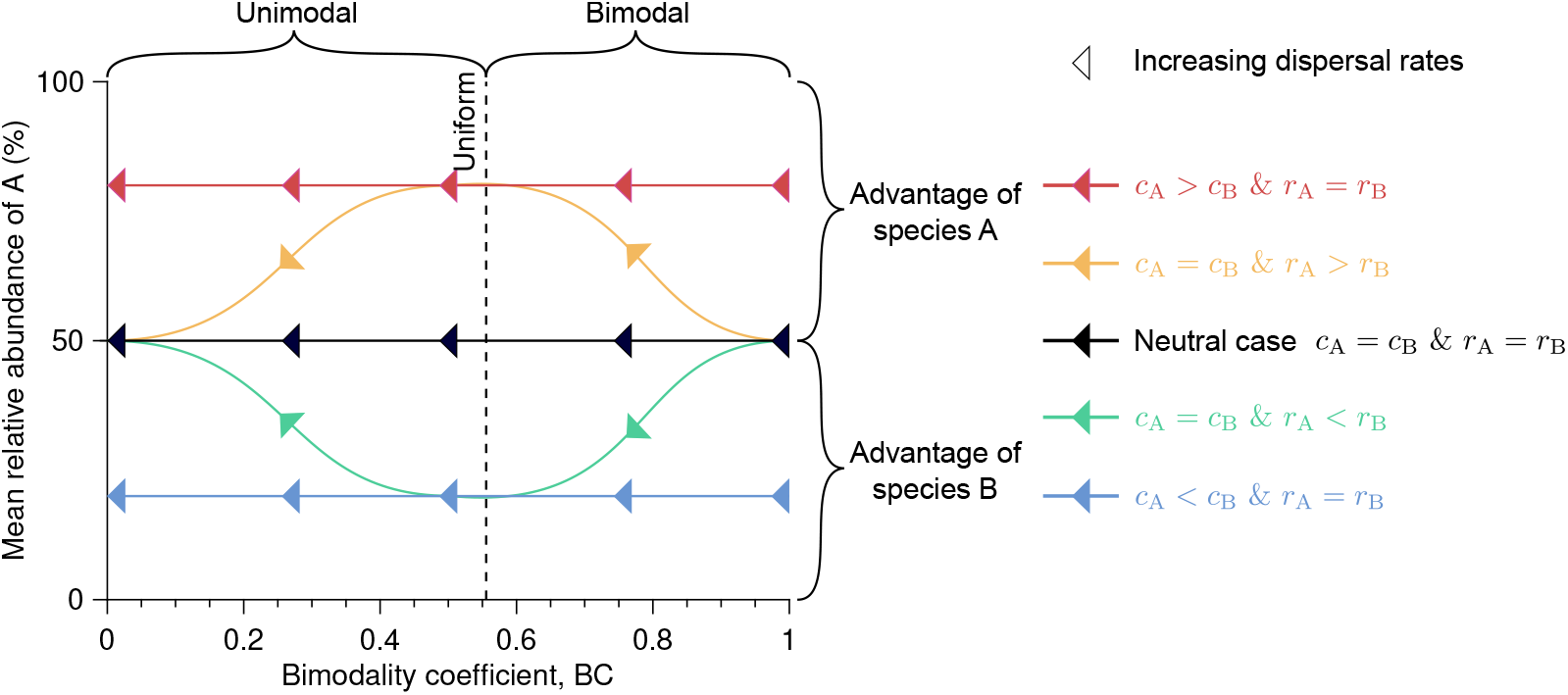
Bimodality coefficient and mean relative abundance characterize how microbial traits compare. This sketch provides a summary of our model predictions. In particular, distribution patterns characterize underlying microbial traits. We consider here the minimal case of two microbial species, namely A and B. Their dispersal rates are denoted by *c*_A_ and *c*_B_, respectively, whereas their division rates are denoted by *r*_A_ and *r*_B_, respectively.

We applied the above reasoning to data collected by Vega and Gore (2017), who populated the gut of *C. elegans* worms by feeding them on a 50/50 mixture of *E. coli* bacteria fluorescently labeled with YFP or dsRed. The authors studied several mixture densities, which allowed them to quantify the microbial community structure across a range of dispersal rates.

Figures 6A shows the mean relative abundance and bimodality coefficients obtained for this data set. As expected, the bimodality coefficient decreases as the bacterial density, and thus the dispersal rates, increases. This decrease reflects the transition from a bimodal to an unimodal relative abundance distribution. The bimodality coefficient is equal to 5/9, which corresponds to a uniform distribution, when the bacterial density is 10^8^ CFU/ml. This bacterial density likely corresponds to a case in which the timescales associated with dispersal and (intra-host) division are similar. Although both strains are expected to neutrally compete, Figure 6A suggests that the dsRed-labeled strain has a consistently higher dispersal rate than the YFP-labeled strain (see Figure 5). To ensure that this conclusion does not result from stochasticity due to the low number of hosts (i.e., 59), we carried out numerical simulations considering two cases. The first case corresponds to the neutral case, where each strain has the same division and dispersal rates, which could legitimately be expected here since the two strains differ only in their labels (i.e., dsRed and YFP). The second case corresponds to one in which one of the strains has a higher dispersal rate than the other but the same division rate, as Figure 6A suggests. As in Vega and Gore (2017)’s experiment, we considered 59 hosts and four different dispersal rates intended to represent different bacterial densities. Figures 6A and B validate that the dispersal rate of dsRed-labeled *E. coli* bacteria is higher than that of YFP-labeled *E. coli* bacteria. It is possible that the strain labeled with dsRed has a better ability to populate a microbiota after dispersal, or that despite the careful setup of the experiment in Vega and Gore (2017), it was present at greater relative abundance in the microbial pool.

**Figure 6:**
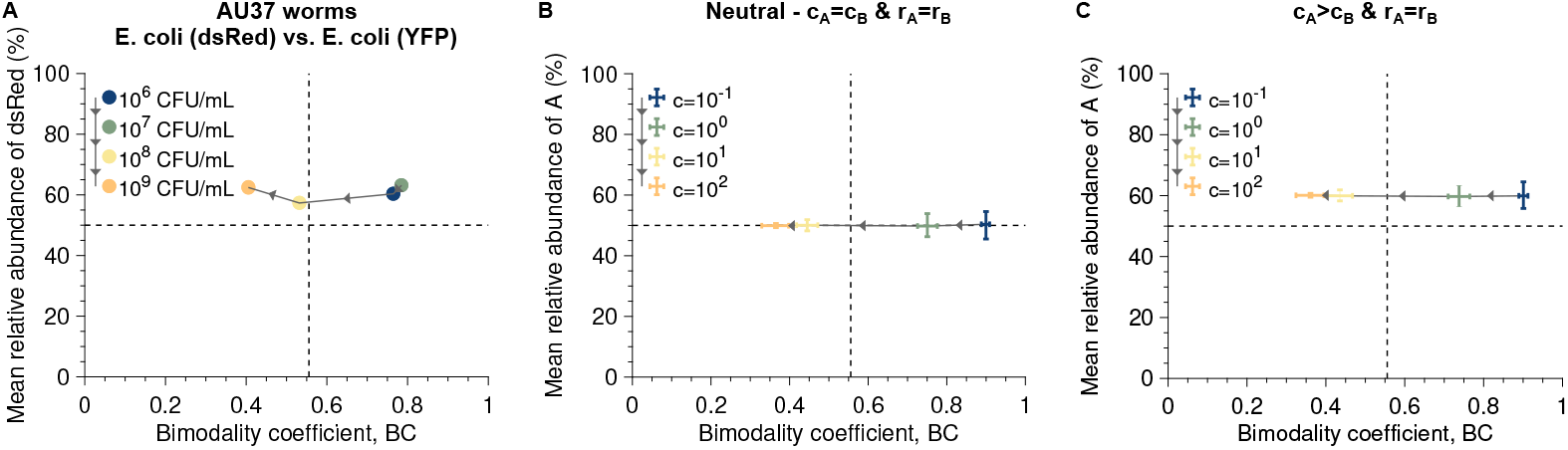
Bimodality coefficient and mean relative abundance indicate a higher dispersal rate in *E. coli* (dsRed) than in *E. coli* (YFP). Sub-figure **A** shows the mean relative abundance of *E. coli* (dsRed) versus the bimodality coefficient for a data set collected by Vega and Gore (2017). Sub-figures **B** and **C** show simulated data for the neutral case and the case where a strain has a higher dispersal rate but the same division rate than the other (i.e., *c*_A_ = 1.5*c*_B_ and *r*_A_ = *r*_B_), respectively. Each data point shows the median calculated from 10^3^ stochastic replicates of the experiment, each with 59 hosts, whereas the bars correspond to the first and third quartiles. Comparing this figure to Figure 5 suggests that *E. coli* (dsRed) has a higher dispersal rate than *E. coli* (YFP). Parameter values: division rates *r*_A_ = *r*_B_ = 5, carrying capacity *K* = 2 × 10^5^.

### Our predictions remain valid at non-zero death rates and with multiple species

A first extension to our model is to include more than two microbial species, as it is relevant to some experimental studies [e.g., Jones et al. (2022)] and real microbial communities. In the Supplementary Material, Section 4, we extended some of our analytical derivations to the case of *S* species. For example, in the neutral case, we show that the low dispersal regime leads to low diversity within communities and high diversity between communities such that each community is populated with only one species, proportionally to its dispersal rate (see Equation S14). Moroever, we demonstrate that the high dispersal regime leads to high diversity within communities and low diversity between communities such that the relative abundances follow a multinomial distribution (see Equation S16). Therefore, eour results on the different dispersal regimes remain valid when including *S* species. We also derive an equation giving the probability that at least two microbial species coexist (see Equation S12, Figure S5).

A second extension to our model is to include death rates, which account for the natural death of microbes and potentially their ejection from the hosts when considering host-associated microbial communities. Including a death rate is expected to change at least two things. First, the final community size no longer equals the carrying capacity, but rather an equilibrium size, which, under a logistic growth, satisfies *K*(1 − *d/r*), where *d* is the death rate. Second, the timescale for dispersal is elongated, as the first disperser survives in the community with probability *d/r*. However, we expect the impact of death to be minimal if the death rate is much lower than the division rate (i.e., *r* ≫ *d*). To validate this prediction, we carried out additional numerical simulations for the neutral case (i.e., *c*_A_ = *c*_B_ = *c* and *r*_A_ = *r*_B_ = *r*) including a death rate *d* that is much lower than the division rate *r* (i.e., *r* ≫ *d*; see the Supplementary Material, Section 5). As shown in Figure S6, our analytical predictions remain valid. The case in which the death rate is larger than the division rate (i.e., *r < d*) would lead the community to decline until it goes extinct, whereas the case in which both rates are similar [i.e., *r* ∼ *d*; see, e.g., Vega and Gore (2017)] is difficult to predict.

## Discussion

### Summary of results

Microbial communities exhibit diversity at multiple scales, whether within and between communities. Recent experiments have shown that this diversity may emerge during the assembly of new communities [e.g., Vega and Gore (2017)]. To deepen our understanding of these experimental results, we built a stochastic model to quantify how dispersal and division contribute to the emergence of diversity during community assembly (see Figure 1). First, we numerically and analytically identified distinct assembly regimes leading to different diversity patterns. These assembly regimes depend on the timescales associated with microbial dispersal and division (see Figure 2). Specifically, if the timescale for dispersal is longer than the timescale for division, i.e., when division is faster than dispersal, each community is populated with only one species, leading to low diversity within communities but high diversity between communities (see Figure 2A). Conversely, if the timescale for dispersal is shorter than the timescale for division, i.e., when division is slower than dispersal, most microbial communities are populated with more than one species, leading to high diversity within communities but low diversity between communities (see Figure 2C). Second, we showed that differences in dispersal rates induce asymmetric relative abundance distributions independent of the assembly regime, such that some species are present at larger abundances than others (see Figure 3). Conversely, differences in division rates appear only in the assembly regime in which the division and dispersal rates are similar (see Figure 4). Our investigation pinpoints two key metrics describing microbial community assembly. Firstly, the bimodality coefficient, which quantifies the number of peaks in relative abundance distribution, determines the assembly regime and, thus, how the dispersal and division rates compare. Secondly, the mean relative abundance describes whether the traits (i.e., the dispersal and division rates) differ across species. Together, these two metrics quantify how the traits of compare across species (see Figure 5). To demonstrate their potential application, we re-analyzed experimental data collected by (Vega and Gore, 2017) (see Figure 6). Overall, our work contributes to a better understanding of microbiome formation by showing how stochasticity, timescales, and microbial traits jointly shape the diversity within and between communities during the assembly of a new microbial community.

### Stochasticity and timescales drive microbial community assembly

The role of stochastic versus deterministic processes in microbial community assembly, and thus in fostering diversity within and between communities, is a fundamental question in microbial ecology (Zhou and Ning, 2017). Experimental studies have shown that isogenic hosts exposed to the same microbial pool can develop different microbiota structures, resulting in diversity within and between communities (Vega and Gore, 2017; Jones et al., 2022). For instance, Vega and Gore (2017) observed considerable heterogeneity in the gut microbiota of *C. elegans* worms fed on the same microbial pool, whereas Jones et al. (2022) found similar patterns in *Drosophila melanogaster* flies. Likewise, human twins exhibit microbiota diversity even within pairs, regardless of zygosity (Goodrich et al., 2014; Vilchez-Vargas et al., 2022). Our stochastic model captures this inherent heterogeneity by modeling microbial dispersal and division as random events. This stochasticity is expected to be relevant whenever a new microbial community is seeded by few individuals initially, independent of its final size.

Our analytical and numerical results reveal distinct assembly regimes. In the *low-dispersal regime*, community assembly is dispersal-limited, leading to greater dissimilarity among communities (i.e., high *β*-diversity) (Etienne and Olff, 2004). Conversely, in the *high-dispersal regime*, dispersal homogenizes community structures (i.e., low *β*-diversity) (Etienne and Olff, 2004). These results echo earlier work. For instance, Zapién-Campos et al. (2020) theoretically investigated the impact of finite host lifespan on the assembly of new microbial communities. They also identified distinct assembly regimes, whose transition from one regime to another depends on a parameter that controls how quickly an unoccupied space is filled with microbes, which is equivalent to the dispersal rate in our model. Our approach extends their work by analytically describing the transition between the different regimes.

Specifically, we built on Vega and Gore (2017)’s study by deriving analytical predictions for the bimodality coefficient, which quantifies the roles of dispersal and division in community assembly. Moreover, we introduced a second metric, the mean relative abundance, which quantifies differences in microbial traits. Reanalyzing Vega and Gore (2017)’s data with our model and these two metrics suggested a different dispersal rate across strains, potentially due to distinct abundances in microbial pool or host colonization efficiency. Importantly, our results remain valid when some assumptions of our model are relaxed. They apply with nonzero microbial death rates, stemming from excretion or death, and extend to communities with more than two microbial species (see the Supplementary Material, Sections 4 and 5). This adaptability highlights our model’s utility for experimental follow-up. Our work thus promises to help identifying patterns and testing hypotheses regarding the forces that drive microbial community assembly.

### Future prospects

Inter-specific interactions occur in our model through density-dependence, which reduces *per capita* division rates as the community size increases. Our model does not take into account more complex inter-specific interactions (Faust and Raes, 2012). For example, some species could benefit from others without either harming or benefiting the others (i.e., commensalism). Since inter-specific interactions are believed to play a key role in shaping microbial communities (Ortiz et al., 2021; Jones et al., 2022), quantifying their impact on the assembly of microbial communities is of great interest (e.g., predation, mutualism, parasitism). In future work, the current model could serve as a null model to test whether more complex interactions are necessary to explain experimental patterns.

In addition to dispersal and selection, mutation plays a crucial role in shaping microbial communities as it generates new genetic diversity (Zhou and Ning, 2017). Mutations are likely to occur in large communities (Vellend, 2010; Nemergut et al., 2013) and during the exponential growth phase associated with community assembly. Yet, the impact of mutation on community assembly has been overlooked so far (Leibold et al., 2004; Chase and Myers, 2011; Stegen et al., 2013; Morlon, 2014), whereas it is necessary to study, for example, the evolution of antibiotic resistance, a pressing public health issue (Baron et al., 2018). The administration of antibiotics to infants during the critical period of microbiota assembly raises concerns for two reasons: it likely reduces intra-host microbial diversity, which is generally associated with host health, and it may promote the establishment of antibiotic-resistant microbes (Reyman et al., 2022). In future work, generating genetic diversity through a mutation probability rate incorporated into the current model would allow us to investigate whether antibiotic resistance emerges during community assembly following a course of antibiotics.

## Author Contributions

LM designed the study; LM performed the numerical and analytical work; LM and CB analyzed and interpreted the data; LM wrote the manuscript; LM and CB edited the manuscript.

## Acknowledgments

The authors thank the THEE Group and Giulia Capella at UniBe for insightful discussion.

