## Supplementary Material for "Drivers of diversity within and between microbial communities during stochastic assembly"

#### Contents

**1 Deterministic description of community assembly** **2**

**2 A stochastic formalism captures additional features of microbial community assembly that complement those described by a deterministic approach** **2**

**3 Very low and high dispersal regimes** **4**

**4 Extension of the model to $S$ species** **5**
- 4.1 Deterministic description of community assembly . . . . . 5
- 4.2 Stochastic description of community assembly . . . . . 6

**5 Extension of the model to death rates** **7**

**6 $\alpha$-diversity** **8**

**7 $\beta$-diversity** **9**

### 1 Deterministic description of community assembly

We begin by establishing expectations for the case of neutral community assembly and for the limiting case of high dispersal in a deterministic model, i.e., by neglecting stochasticity.

**System of ODEs.** We describe the dynamics of the community assembly by a system of ordinary differential equations (ODEs)

$$\begin{cases} \frac{\mathrm{d}N_\mathrm{A}}{\mathrm{d}t} = (c_\mathrm{A} + r_\mathrm{A} N_\mathrm{A}) \left(1 - \frac{N_\mathrm{A}+N_\mathrm{B}}{K}\right) , \\ \frac{\mathrm{d}N_\mathrm{B}}{\mathrm{d}t} = (c_\mathrm{B} + r_\mathrm{B} N_\mathrm{B}) \left(1 - \frac{N_\mathrm{A}+N_\mathrm{B}}{K}\right) , \end{cases} \tag{S1}$$

with the initial conditions $N_\mathrm{A} = N_\mathrm{B} = 0$ since we consider initially microbe-free microbial communities. We rewrite the system 1 as

$$\begin{cases} \frac{\mathrm{d}N_\mathrm{A}}{c_\mathrm{A}+r_\mathrm{A}N_\mathrm{A}} = \left(1 - \frac{N_\mathrm{A}+N_\mathrm{B}}{K}\right) \mathrm{d}t \, , \\ \frac{\mathrm{d}N_\mathrm{B}}{c_\mathrm{B}+r_\mathrm{B}N_\mathrm{B}} = \left(1 - \frac{N_\mathrm{A}+N_\mathrm{B}}{K}\right) \mathrm{d}t \, , \end{cases} \tag{S2}$$

the solution of which reads

$$\left(1 + \frac{r_\mathrm{A}}{c_\mathrm{A}} N_\mathrm{A}\right)^{\frac{1}{r_\mathrm{A}}} = \left(1 + \frac{r_\mathrm{B}}{c_\mathrm{B}} N_\mathrm{B}\right)^{\frac{1}{r_\mathrm{B}}} . \tag{S3}$$

**Stationary solution.** To determine whether the two microbial species A and B coexist at the end of the community assembly, one can quantify the stationary solution from Equation 3 combined with $N_\mathrm{A} + N_\mathrm{B} = K$. In the neutral case, in which both microbial species have the same dispersal and division rates (i.e., $c_\mathrm{A} = c_\mathrm{B} = c$ and $r_\mathrm{A} = r_\mathrm{B} = r$), we obtain

$$N_\mathrm{A} = N_\mathrm{B} = \frac{K}{2} . \tag{S4}$$

Thus, in the neutral case, the deterministic model yields that both microbial species are expected to coexist at equal abundances at the end of the community assembly. A second limiting case involves microbial species whose dispersak rates are much greater than their division rates (i.e., $c_\mathrm{A} \gg r_\mathrm{A}$ and $c_\mathrm{B} \gg r_\mathrm{B}$), which leads to

$$N_\mathrm{A} = \frac{c_\mathrm{A}}{c_\mathrm{A} + c_\mathrm{B}} K \text{ and } N_\mathrm{B} = \frac{c_\mathrm{B}}{c_\mathrm{A} + c_\mathrm{B}} K . \tag{S5}$$

Thus, when dispersal is much faster than division, the community assembly is predicted to be fully driven by dispersal. Specifically, the greater the dispersal rate of a microbial species, the larger the expected abundance of this species in the final structure of the microbial community.

### 2 A stochastic formalism captures additional features of microbial community assembly that complement those described by a deterministic approach

There are two main types of mathematical models to describe community assembly. Deterministic models are based on systems of ordinary differential equations describing the dynamics of the community and population sizes (see Equation 1), whereas stochastic models are based on master equations describing the dynamics of the probability of the community and population sizes. Since microbial communities involve a large number of individuals, one could argue that

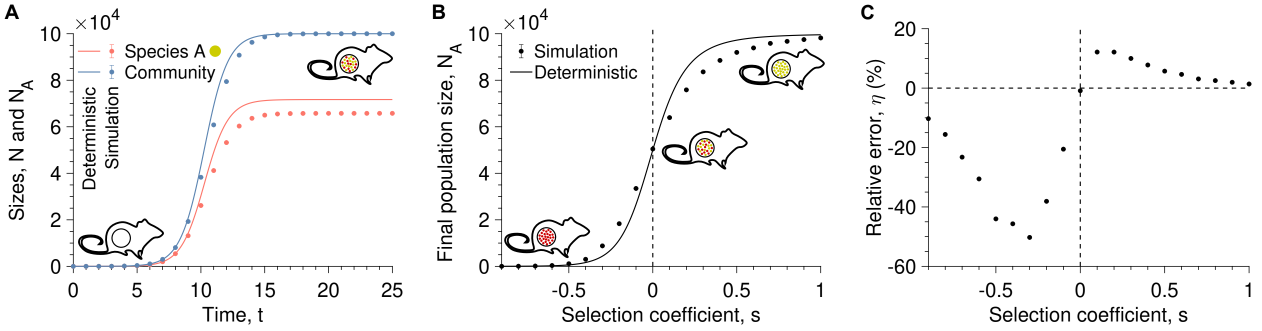


Figure S1: **Stochasticity counteracts selection in community assembly, leading deterministic models to underestimate selection.** Panel **A** shows the community size and number of A microbes (i.e., $N = N_\mathrm{A} + N_\mathrm{B}$ and $N_\mathrm{A}$, respectively) as a function of time $t$ with a selection coefficient of A microbes equal to 0.1 (i.e., $r_\mathrm{A} = 1.1$ and $r_\mathrm{B} = 1$, leading to $s = r_\mathrm{A} - r_\mathrm{B} = 0.1$). Panel **B** shows the final number of A microbes once the carrying capacity is reached (i.e., $N_\mathrm{A} + N_\mathrm{B} = K$) versus the selection coefficient $s = r_\mathrm{A} - r_\mathrm{B}$. Panel **C** shows the relative error $\eta$ between the simulated data and deterministic predictions from Sub-Figure **B** against the selection coefficient $s$. In panels **A** and **B**, the solid lines represent analytical predictions (see Equation 1), and markers correspond to simulated data averaged over $10^3$ stochastic replicates. In panel **C**, the vertical dashed line shows the neutral case (i.e., $r_\mathrm{A} = r_\mathrm{B}$, leading to $s = 0$). Parameter values: carrying capacity $K = 10^5$, dispersal rates $c_\mathrm{A} = c_\mathrm{B} = 1$.

a deterministic approach would be a relevant choice. However, when it comes to community assembly, the process initially involves small community and population sizes, which are conducive to stochastic demography. We, therefore, check here how the deterministic approach describes the community assembly dynamics compared to its stochastic counterpart.

Figure S1A shows the mean community and population sizes during the community assembly, averaged over several stochastic replicates, in a case in which both microbial species disperse at the same rate (i.e., $c_\mathrm{A} = c_\mathrm{B}$), whereas A microbes reproduce slightly more often than B microbes (i.e., $r_\mathrm{A} = 1.1$ and $r_\mathrm{B} = 1$, leading to $s = 0.1$). In the growth phase, both the mean community size and number of A microbes are overestimated by the deterministic equations (see Equation 1). Although the final community size, i.e., once the carrying capacity is reached, is well predicted by the deterministic equations, the final number of A microbes is overestimated.

Figure S1B shows the final number of A microbes for different selection coefficients $s$. Except in the neutral case in which both microbial species have the same division rate (i.e., $r_\mathrm{A} = r_\mathrm{B}$, leading to $s = 0$), the deterministic approach poorly predicts the final number of A microbes. More specifically, if the selection coefficient is negative (i.e., $s < 0$), it underestimates the final number of A microbes, whereas if the selection coefficient is positive (i.e., $s > 0$), it overestimates the final number of A microbes. As shown in Figure S1C, the relative error $\eta$, which quantifies the difference between the simulated data and the deterministic predictions of Figure S1B, ranges from -50% to 12%. Thus, the deterministic approach leads to poor predictions of the community assembly.

As explained by Marrec et al. (2023), the discrepancy between simulated data and deterministic predictions results from the variability of the waiting times between dispersal and division events. This variability, which is particularly important at small initial community and population sizes and, thus, occurs independently of the carrying capacity, is not taken into account in deterministic approaches. These results highlight the importance of using a stochastic formalism when studying microbial community assembly. As we will show in the following, demographic stochasticity is not the only reason why a stochastic description is more suitable for describing the dynamics of microbial community assembly than a deterministic description.

### 3 Very low and high dispersal regimes

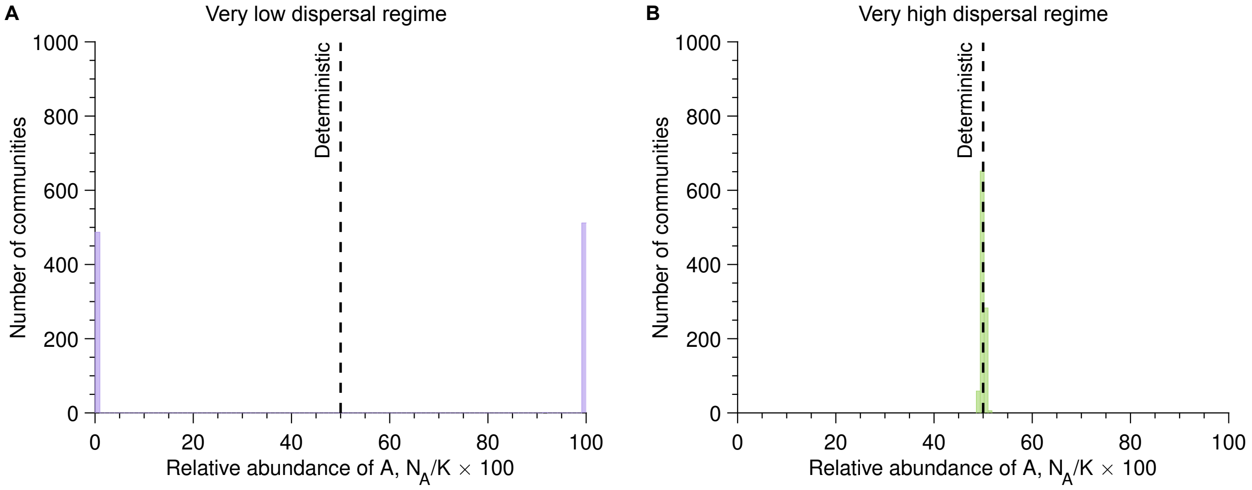


Figure S2: **Very low and high dispersal regimes lead to bimodal and unimodal species abundance distributions, respectively.** Sub-figures **A** and **B** show the number of communities versus the percentage of A microbes once the carrying capacity is reached (i.e., $N_A/K \times 100$) with the dispersal rate equal to $c = 10^{-4}$ and $c = 10^4$, respectively (here $c_A = c_B = c$). Parameter values: division rates $r_A = r_B = 1$, carrying capacity $K = 10^5$, number of hosts $10^3$.

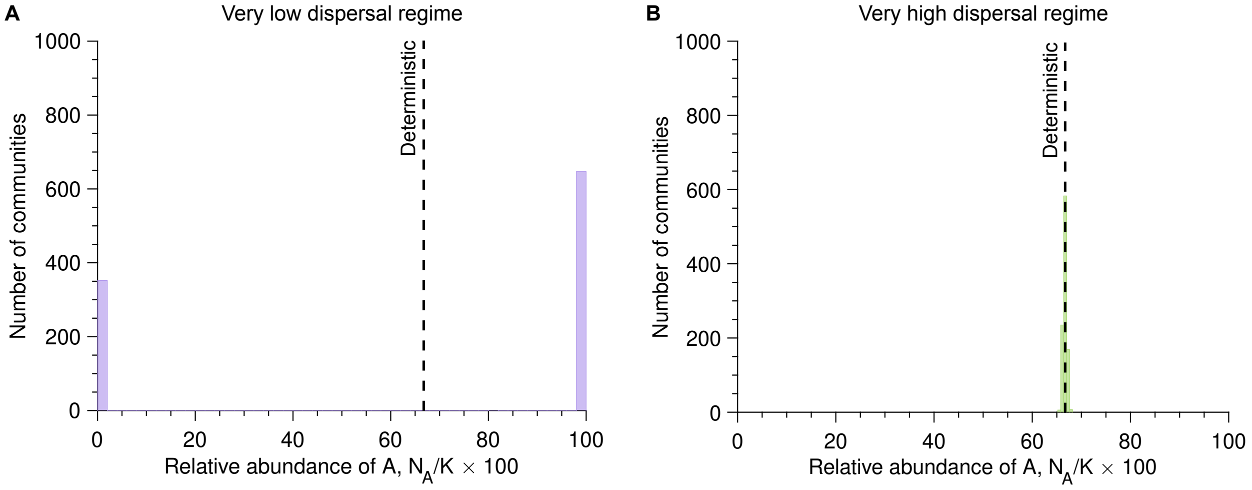


Figure S3: **Differences in dispersal rates make species abundance distributions asymmetric.** Sub-figures **A** and **B** show the number of communities versus the percentage of A microbes once the carrying capacity is reached (i.e., $N_A/K \times 100$) with the dispersal rate equal to $c = 10^{-4}$ and $c = 10^4$, respectively (here $c_A = 2c_B$). Parameter values: division rates $r_A = r_B = 1$, carrying capacity $K = 10^5$, number of communities $10^3$.

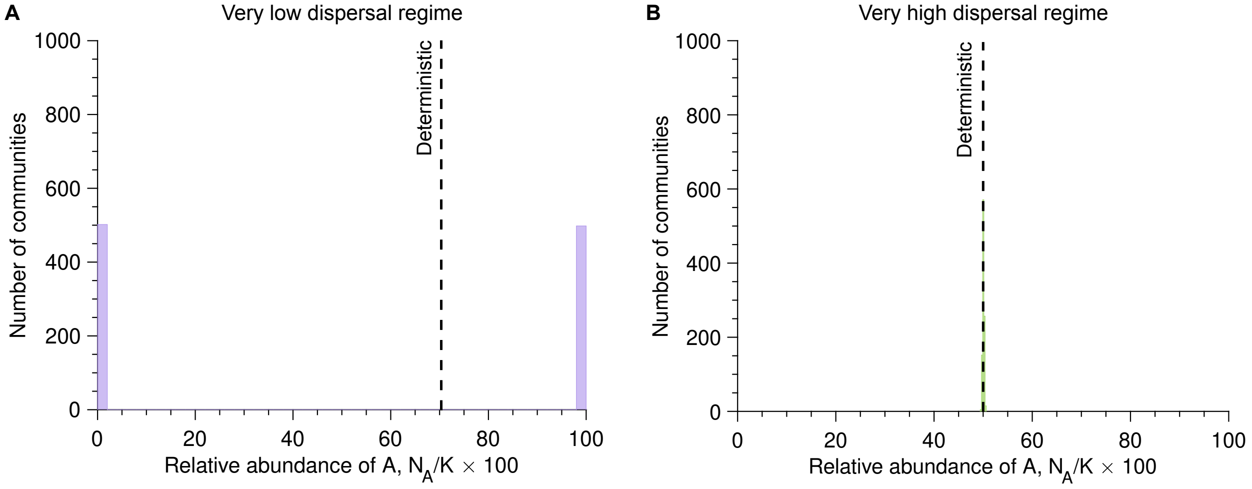


Figure S4: **Selection does not appear in the very high and very low dispersal regimes.** Sub-figures **A** and **B** show the number of communities versus the percentage of A microbes once the carrying capacity is reached (i.e., $N_{\mathrm{A}}/K \times 100$) with the dispersal rate equal to $c = 10^{-4}$ and $c = 5 \times 10^5$, respectively (here $c_{\mathrm{A}} = c_{\mathrm{B}}$). Parameter values: division rate of A microbes $r_{\mathrm{A}} = 1.05$, division rate of B microbes $r_{\mathrm{B}} = 1$, selection coefficient $s = 0.05$, carrying capacity $K = 10^5$, number of communities $10^3$.

### 4 Extension of the model to $S$ species

In this section, we extend our model from 2 to $S$ species. The only difference with the model presented in the main text is that the microbial pool is fixed and composed of $S$ species, denoted by $i = 1, 2, ..., S$. Their division rates are denoted $r_i$ whereas their dispersal rates are denoted by $c_i$.

#### 4.1 Deterministic description of community assembly

**System of ODEs.** The system of ordinary differential equations (ODEs) describing the community assembly is given by

$$\frac{\mathrm{d}N_i}{\mathrm{d}t} = (c_i + r_i N_i)\left(1 - \frac{\sum_{j=1}^{S} N_j}{K}\right) \text{ with } 1 \le i \le S\,. \tag{S6}$$

**Stationary solution.** To determine whether the $S$ species coexist at the end of the community assembly, one can quantify the stationary solution from Equation S6 combined with $\sum_{i=1}^{S} N_i = K$. In the neutral case, in which the $S$ species have the same dispersal and division rates (i.e., $c_i = c$ and $r_i = r$ with $1 \le i \le S$), one obtains

$$N_i = \frac{K}{S} \text{ with } 1 \le i \le S\,. \tag{S7}$$

Thus, in the neutral case, the $S$ species are expected to coexist in equal abundances at the end of the community assembly. A second interesting case involves species whose dispersal rates are much greater than their division rates (i.e., $c_i \gg r_i$ with $1 \le i \le S$), which leads to

$$N_i = \frac{c_i}{\sum_{j=1}^{S} c_j} K \text{ with } 1 \le i \le S\,. \tag{S8}$$

Thus, when the timescale associated with dispersal is much shorter than that associated with division, the community assembly is fully driven by dispersal. Specifically, the greater the dispersal rate of a species, the larger the abundance of this species in the final structure of the microbial community.

#### 4.2 Stochastic description of community assembly

The system of equations governing the dynamics of the probability $P(\{N_1, N_2, ..., N_S\}, N)$ that a community of size $N$ has a structure $\{N_1, N_2, ..., N_S\}$, where $\sum_{i=1}^{S} N_i = N$, satisfies (Gardiner, 2009; Van Kampen, 2011)

$$\begin{aligned} P(\{N_1, N_2, ..., N_S\}, N+1) = & \sum_{j=1}^{S} \alpha_N^{N_j - 1} P(\{N_1, N_2, ..., N_j - 1, ..., N_S\}, N) \\ & + (1 - \sum_{j=1}^{S} \alpha_N^{N_j}) P(\{N_1, N_2, ..., N_S\}, N)\,, \end{aligned} \tag{S9}$$

where $\alpha_N^{N_i}$ is the probability that the increase in community size by one individual (i.e., $N \to N+1$) is due to the dispersal or division of a microbe of species $i$. This probability is given by

$$\alpha_N^{N_i} = \frac{r_i N_i + c_i}{\sum_{j=1}^{S} (r_j N_j + c_j)}\,. \tag{S10}$$

Since we consider in our model that the microbial assembly starts from a microbe-free community, the master equation SS9 admits as initial condition $P(\{0, 0, ..., 0\}, 0) = 1$. One can compute the probability $P_i$ that the community is populated with only microbes of species $i$, which is given by

$$P_i = \frac{c_i}{\sum_{j=1}^{S} c_j} \frac{\left(1 + \frac{c_i}{r_i}\right)_{K-1}}{\left(1 + \frac{\sum_{j=1}^{S} c_j}{r_i}\right)_{K-1}}\,. \tag{S11}$$

The previous equation allows us to determine whether coexistence of the $S$ species occurs at the end of the assembly process. More specifically, the probability that the $S$ species coexist, denoted by $P_{\text{coexist}}$, is given by

$$P_{\text{coexist}} = 1 - \sum_{i=1}^{S} P_i = 1 - \sum_{i=1}^{S} \frac{c_i}{\sum_{j=1}^{S} c_j} \frac{\left(1 + \frac{c_i}{r_i}\right)_{K-1}}{\left(1 + \frac{\sum_{j=1}^{S} c_j}{r_i}\right)_{K-1}}\,. \tag{S12}$$

An example is shown in Figure S5.

**Low dispersal regime.** In the case in which the dispersal rates are much lower than the division rates (i.e., $c_i \ll r_i$ with $1 \leq i \leq S$), the probability $\alpha_N^{N_i}$ (see Equation S10) can be simplified into

$$\alpha_N^{N_i} = \frac{r_i N_i}{\sum_{j=1}^{S} r_j N_j}\,. \tag{S13}$$

In addition, Equation S11 can be simplified into

$$P_i \underset{c_i \ll r_i}{\approx} \frac{c_i}{\sum_{j=1}^{S} c_j}\,. \tag{S14}$$

**High dispersal regime.** In the case in which the dispersal rates are much higher than the division rates (i.e., $c_i \gg r_i$ with $1 \leq i \leq S$), the probability $\alpha_N^{N_i}$ (see Equation S10) can be simplified into

$$\alpha_N^{N_i} = \frac{c_i}{\sum_{j=1}^{S} c_j} = \alpha_i \, . \tag{S15}$$

In this case, the species abundance follows a multinomial distribution, which satisfies

$$P(\{N_1 = n_1, N_2 = n_2, ..., N_S = n_S\}, N) = \frac{N!}{n_1! n_2! ... n_S!} \alpha_1^{n_1} \alpha_2^{n_2} ... \alpha_S^{n_S} \, . \tag{S16}$$

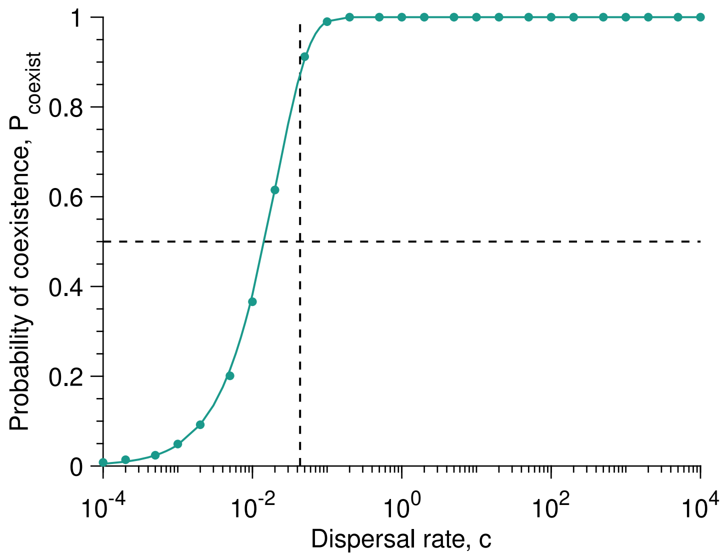


Figure S5: **Our analytical derivations can be extended to more than two species.** Probability of coexistence $P_{\text{coexist}}$ versus the dispersal rate $c$ with $S = 5$ species in the microbial pool assuming the neutral case (i.e., $c_i = c$ and $r_i = r$ for $1 \leq i \leq S$). The data points are averaged over $10^3$ replicates (or communities), whereas the solid line represents our analytical prediction (see Equation S12). Parameter values: division rate $r = 1$, carrying capacity $K = 10^5$.

### 5 Extension of the model to death rates

In this section, we extend our model so that it accounts for death rates. Specifically, the death rate of A microbes is denoted by $d_{\text{A}}$, whereas the death rate of B microbes is denoted by $d_{\text{B}}$.

**System of ODEs.** When including death rates, the system of ordinary differential equations (ODEs) describing the dynamincs of community assembly is given by

$$\begin{cases} \frac{\mathrm{d}N_{\text{A}}}{\mathrm{d}t} = (c_{\text{A}} + r_{\text{A}} N_{\text{A}}) \left(1 - \frac{N_{\text{A}}+N_{\text{B}}}{K}\right) - d_{\text{A}} N_{\text{A}} \, , \\ \frac{\mathrm{d}N_{\text{B}}}{\mathrm{d}t} = (c_{\text{B}} + r_{\text{B}} N_{\text{B}}) \left(1 - \frac{N_{\text{A}}+N_{\text{B}}}{K}\right) - d_{\text{B}} N_{\text{B}} \, . \end{cases} \tag{S17}$$

**Gillespie algorithm simulating the microbial community assembly.** In our model including death, we have six distinct events, among which the divisions of the microbes of species A and B

$$A \xrightarrow{r_{\text{A}}\left(1 - \frac{N_{\text{A}}+N_{\text{B}}}{K}\right) N_{\text{A}}} 2A \, ,$$

$$B \xrightarrow{r_\mathrm{B}\left(1-\frac{N_\mathrm{A}+N_\mathrm{B}}{K}\right)N_\mathrm{B}} 2B\,,$$

the dispersal of the microbes of species A and B

$$\emptyset \xrightarrow{c_\mathrm{A}\left(1-\frac{N_\mathrm{A}+N_\mathrm{B}}{K}\right)} A\,,$$

$$\emptyset \xrightarrow{c_\mathrm{B}\left(1-\frac{N_\mathrm{A}+N_\mathrm{B}}{K}\right)} B\,,$$

and the death of the microbes of species A and B

$$A \xrightarrow{d_\mathrm{A}N_\mathrm{A}} \emptyset\,,$$

$$B \xrightarrow{d_\mathrm{B}N_\mathrm{B}} \emptyset\,,$$

Note that the community size $N = N_\mathrm{A}+N_\mathrm{B}$ can decrease when including death . The simulation steps are as follows:

1. Initialization: The microbial community starts from $N_\mathrm{A} = 0$ and $N_\mathrm{B} = 0$ microbe at time $t = 0$.

2. Time update: The time increment $\Delta t$ is randomly sampled from an exponential distribution with mean $1/\left((r_\mathrm{A}N_\mathrm{A} + c_\mathrm{A} + r_\mathrm{B}N_\mathrm{B} + c_\mathrm{B})(1 - (N_\mathrm{A} + N_\mathrm{B})/K) + d_\mathrm{A}N_\mathrm{A} + d_\mathrm{B}N_\mathrm{B}\right)$ and the time is updated by $t \leftarrow t + \Delta t$.

3. Event selection: The next event to occur is chosen randomly proportionally to its probability.

4. Population size update: The population sizes are updated according to the event selected in Step 3. For example, if the division of a species A microbe is chosen, the population size of species A is updated by $N_\mathrm{A} \leftarrow N_\mathrm{A} + 1$.

5. We return to Step 2 until the community size reaches for the first time the value defined by the user.

We consider that each stochastic realization of the above algorithm describes the assembly of the microbial community of a single community. Thus, collecting several stochastic realizations is equivalent to simulating the microbial assembly of several communities.

### 6 $\alpha$-diversity

$\alpha$-diversity quantifies the species diversity (i.e., richness) within a community. There exist different ways to calculate it. For example, the Shannon Index quantifies both species richness and evenness, but with weight on the richness, which is given by

$$\mathrm{Shannon} = -\sum_{i=1}^{R} p_i \ln(p_i)\,, \tag{S18}$$

where $p_i$ is the abundance of species $i$ and $R$ the total number of species ($R = 2$ in the main text). The Simpson Index is based on the probability that two microbes taken from the community at random are of different species. As this is a probability, its value ranges from 0 to 1 and is given by

$$\mathrm{Simpson} = 1 - \sum_{i=1}^{R} p_i^2\,. \tag{S19}$$

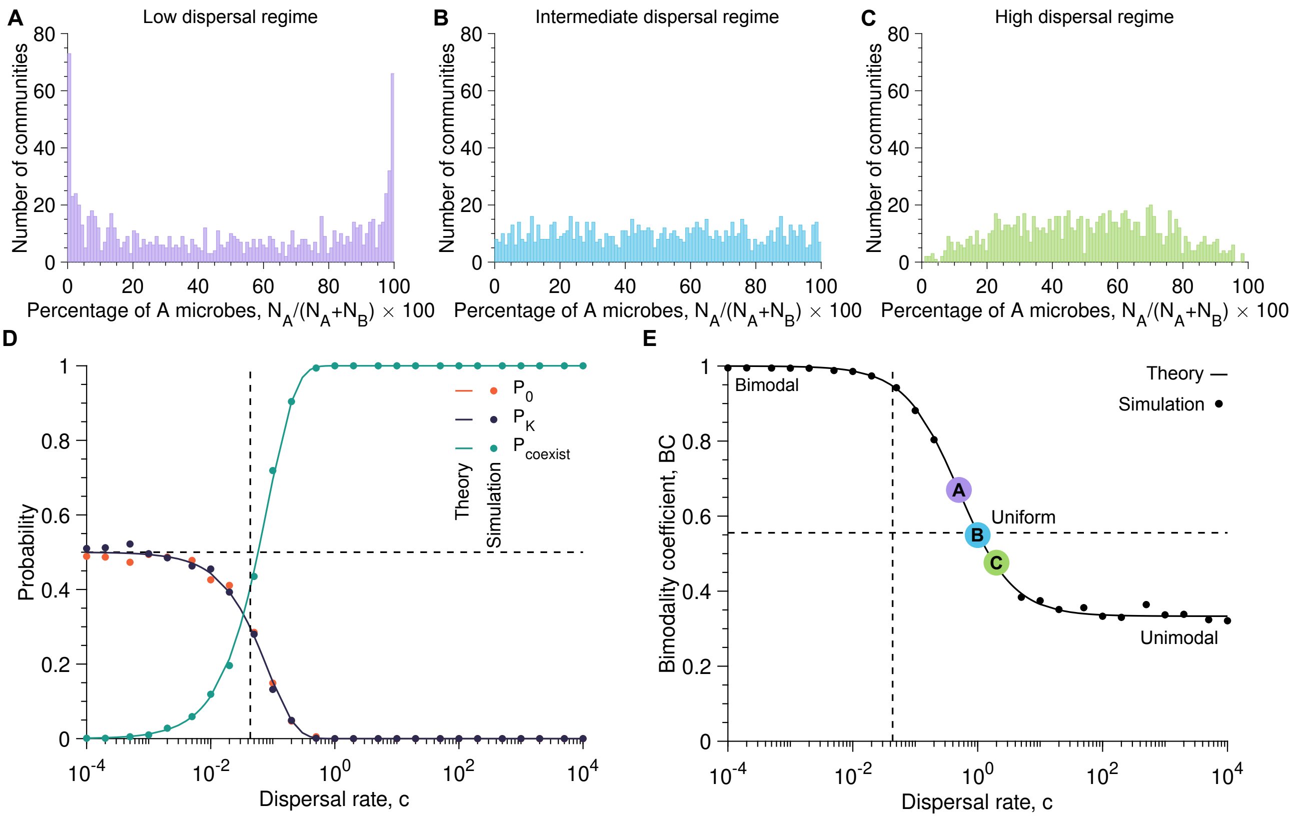


Figure S6: **Our analytical predictions remain valid with a low death rate.** Sub-figures **A**, **B**, and **C** show the number of communities versus the percentage of A microbes once the equilibrium size is reached (i.e., $N_{\mathrm{A}} + N_{\mathrm{B}} = K(1 - d/r)$) with the dispersal rate equal to $c = 0.5$, $c = 1$, and $c = 2$, respectively (here $c_{\mathrm{A}} = c_{\mathrm{B}} = c$). Sub-figure **D** represents the probabilities that, at the end of the microbial community assembly, the community is left with no A microbes (i.e., $P_0$), has only A microbes (i.e., $P_K$), and is composed of both species (i.e., $P_{\mathrm{coexist}} = 1 - P_0 - P_K$) as a function of the dispersal rate $c$. Sub-figure **E** presents the bimodality coefficient BC against the dispersal rate $c$. In Sub-figures **D** and **E**, the solid lines show analytical predictions, whereas the markers correspond to simulated data averaged over $10^3$ stochastic replicates, each considered a community. In Sub-figures **D** and **E**, the vertical dashed line shows the dispersal rate value at which the timescales associated with dispersal and division are equal (i.e., $c = c_{\mathrm{lim}}$). In Sub-figures **D** and **E**, the vertical dashed line shows a probability of 1/2 and a bimodality coefficient of 5/9, respectively. Parameter values: division rates $r_{\mathrm{A}} = r_{\mathrm{B}} = 1$, death rates $d_{\mathrm{A}} = d_{\mathrm{B}} = 0.1$, carrying capacity $K = 10^5$, number of communities $10^3$.

### 7 $\beta$-diversity

$\beta$-diversity quantifies the differences between communities. One of the most common ways to calculate it is to use the Bray-Curtis dissimilarity, which quantifies the abundances of microbes shared between two communities. The Bray-Curtis dissimilarity ranges from 0 to 1 and is given by

$$\mathrm{Bray} - \mathrm{Curtis} = 1 - \frac{2C_{ij}}{S_i + S_j}, \tag{S20}$$

where $C_{ij}$ is the sum of the lowest values for species in common between the two communities, whereas $S_i$ and $S_j$ are the total number of microbes founds in communities $i$ and $j$, respectively (here $S_i = S_j = K$).

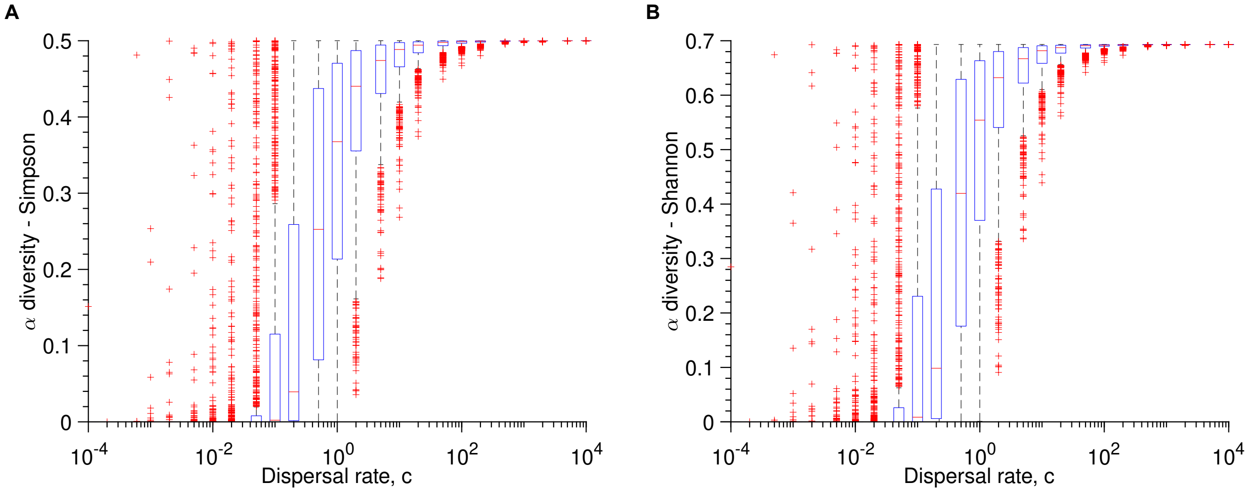


Figure S7: **$\alpha$-diversity increases as dispersal rate increases.** Sub-figures **A** and **B** show the Simpson and Shannon indexes, respectively, as a function of the dispersal rate $c$ (here $c_{\mathrm{A}} = c_{\mathrm{B}} = c$). Parameter values: division rates $r_{\mathrm{A}} = r_{\mathrm{B}} = 1$, carrying capacity $K = 10^5$, number of communities $10^3$.

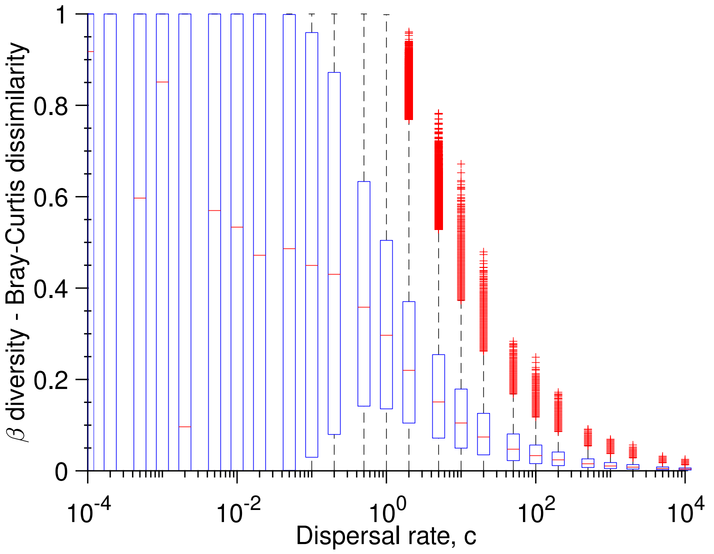


Figure S8: **$\beta$-diversity decreases as dispersal rate increases.** The Bray-Curtis dissimilarity as a function of the dispersal rate $c$ (here $c_{\mathrm{A}} = c_{\mathrm{B}} = c$). Parameter values: division rates $r_{\mathrm{A}} = r_{\mathrm{B}} = 1$, carrying capacity $K = 10^5$, number of communities $10^3$.
